# Tissue microenvironment dictates the state of human induced pluripotent stem cell-derived endothelial cells of distinct developmental origin in 3D cardiac microtissues

**DOI:** 10.1101/2022.11.22.517426

**Authors:** Xu Cao, Maria Mircea, Francijna E. van den Hil, Hailiang Mei, Katrin Neumann, Konstantinos Anastassiadis, Christine L. Mummery, Stefan Semrau, Valeria V. Orlova

**Author notes:** Contributed equally (Xu Cao, Maria Mircea).

## Abstract

Each tissue and organ in the body has its own type of vasculature. Here we demonstrate that organotypic vasculature for the heart can be recreated in a three-dimensional cardiac microtissue (MT) model composed of human induced pluripotent stem cell (hiPSC)-derived cardiomyocytes (CMs), cardiac fibroblasts (CFs) and endothelial cells (ECs). ECs in cardiac MTs upregulated expression of markers enriched in human intramyocardial ECs (iECs), such as *CD36, CLDN5, APLNR, NOTCH4, IGFBP3, ARHGAP18*, which were previously identified in the single-cell RNA-seq dataset from the human fetal heart (6.5-7 weeks post coitum). We further show that the local microenvironment largely dictates the organ-specific identity of hiPSC-derived ECs: we compared ECs of different developmental origins derived from two distinct mesoderm subtypes (cardiac and paraxial mesoderm) and found that independent of whether the ECs were cardiac or paraxial mesoderm derived, they acquired similar identities upon integration into cardiac microtissues. This was confirmed by single-cell RNA-seq. Overall, the results indicated that whilst the initial gene profile of ECs was dictated by developmental origin, this could be modified by the local tissue environment such that the original identity was lost and the organotypic identity acquired through local environmental signals. This developmental “plasticity” in ECs has implications for multiple pathological and disease states.

## INTRODUCTION

Development of the vascular system is one of the earliest events in organogenesis and defects in this process often result in embryonic and postnatal lethality. Endothelial cells (ECs) that form the inner lining of blood and lymphatic vessels are specialized cells that adapt to local microenvironmental cues to support the function of various organs. The heart is a prime example of the importance that interplay between the vasculature and the myocardium has in organ growth, remodeling, and function. ECs in the heart originate from several developmental lineages that later converge to similar states depending on their location. Structurally and functionally, ECs in the heart can be divided into the endocardium, intramyocardial capillary ECs, coronary arteries/veins and lymphatic ECs. During development, heart ECs predominantly originate in the lateral plate mesoderm that includes both pre-cardiac and cardiac mesoderm (Milgrom-Hoffman et al., 2011). Endocardium and *sinus venosus* are the two major sources on intramyocardial and coronary ECs (He and Zhou, 2018; Sharma et al., 2017a, 2017b; Tian and Zhou, 2022). Furthermore, genetic ablation of the *sinus venosus* EC lineage results in compensation from the endocardial EC lineage (Sharma et al., 2017b). Endocardium- and *sinus venosus-derived* EC progenitors, converge to an increasingly similar state that is dictated by the local microenvironment, despite being initially transcriptionally distinct (Phansalkar et al., 2021). In addition, recent studies showed that cardiac lymphatic ECs predominately originate from two lineages, namely Isl1+ second heart field and Pax3+ paraxial mesoderm (PM) (Lioux et al., 2020; Stone and Stainier, 2019), which further increases the spectrum of developmental origins of ECs in the heart.

Endothelial cells in the intramyocardial capillaries, or iECs, constitute a specialized barrier between the blood and the myocardium. They deliver oxygen and essential nutrients, specifically fatty acids, to the cardiomyocytes to fulfill high energy demands of the working myocardium (Brutsaert, 2003). Endocardial ECs (eECs) form the lining of the inner surface in the ventricles and atria and play important roles during the development of working myocardium, such as formation of trabeculae network and compaction of the myocardium (Qu et al., 2022). Analysis of developing mouse heart identified markers that distinguish eECs and iECs (Iso et al., 2018; Zhang et al., 2016a). Recent single-cell RNA sequencing (scRNA-seq) studies of the human fetal heart confirmed that some of the markers identified in mouse are conserved in human, including *CD36* and *FABP5* for iECs and *NPR3* and *CDH11* for eECs, among others (McCracken et al., 2022; Miao et al., 2020).

Human pluripotent stem cells (hPSCs) represent a valuable *in vitro* model to study early stages of human development, including the heart (Hofbauer et al., 2021a). Over the past several years, protocols to differentiate cardiac cell types from hPSCs, such as different sub-types of cardiomyocytes, epicardial cells, cardiac fibroblasts and ECs were developed (Thomas et al., 2022). Engineered multicellular cardiac tissues can be generated by combining these different cell types (Campostrini et al., 2021). These have proved useful to investigate the contribution of non-cardiac cell types to cardiomyocyte cell maturation and disease (Giacomelli et al, 2020a). In addition, methods to create multicellular cardioids from hPSCs have been developed to model embryonic stages from self-organized cardiac and foregut structures to early morphogenesis during heart tube formation (Drakhlis et al., 2021; Hofbauer et al., 2021b; Lewis-Israeli et al., 2021; Silva et al., 2021). Recent studies showed that by following a specialized developmental program, hPSCs can be differentiated into ECs from several developmental lineages, such as extra- and intraembryonic hemogenic ECs (Ditadi et al., 2015; Ng et al., 2016; Uenishi et al., 2014), endocardial ECs (Mikryukov et al., 2021) and liver sinusoidal ECs (Gage et al., 2020). We previously developed a method to co-differentiate cardiomyocytes and ECs from hPSCs from cardiac mesoderm (Giacomelli et al., 2017). These hPSC-derived ECs expressed a number of cardiac specific genes like *MEOX2, GATA4, GATA6* and *ISL1*, while tissue specific endocardial or intramyocardial markers were still absent, likely because of the lack of local microenvironmental cues.

Recently, we established a three-dimensional (3D) cardiac microtissue (MT) model composed of ECs, cardiomyocytes and cardiac fibroblasts, all derived from hiPSCs (Giacomelli et al., 2020). We showed that hPSC-derived cardiomyocytes in cardiac 3D MTs showed enhanced functional and structural maturation via interaction with ECs and cardiac fibroblasts. We further showed that developmental origin of the fibroblasts was critical in this model, as cardiac-, but not skin-, fibroblasts supported cardiomyocyte maturation in 3D MTs.

However, whether 3D cardiac MTs actually induce organ-specific characteristics and to a what extent the developmental origin of ECs plays a role has not been investigated. We therefore aimed here to address these questions by comparing ECs derived from two distinct mesoderm sub-types: MESP1+ cardiac mesoderm and PAX3+ paraxial mesoderm. To do this, we utilized our earlier protocol to differentiate cardiac mesoderm-derived ECs (Giacomelli et al., 2017) and developed a new protocol to differentiate ECs from paraxial mesoderm. Cardiac MTs were then generated using these two sources of ECs. Although newly differentiated ECs from the two origins showed distinct identities, they strikingly became more similar after extended culture in cardiac MTs. Furthermore, based on eEC- and iEC-specific signatures extracted from a published scRNA-seq dataset of human fetal heart (Asp et al., 2019), we observed an iEC rather than an eEC identity for both developmental origins after MT culture. In summary, this study shows that although certain characteristics are inherited from progenitors, ECs are “plastic” and efficiently adapt to the microenvironment to acquire new tissue-specific signatures. Our results provide new insights into how organ/tissue-specific cell identities are acquired; this will inform the preparation of hiPSC -derived, organ specific ECs for disease modeling and drug development but as importantly, will provide a platform for understanding how EC plasticity might be regulated by microenvironmental context.

## RESULTS

### Differentiation of endothelial cells from cardiac and paraxial mesoderm

We set out to derive ECs from human induced pluripotent stem cells (hiPSCs) via both cardiac and paraxial mesoderm intermediates. To obtain ECs from cardiac mesoderm (CMECs), we used a protocol established previously in our group (Giacomelli et al., 2017) (Figure 1A). Briefly, BMP4 (20 ng/ml), Activin A (ACTA, 20 ng/ml) and CHIR99021 (CHIR, 1.5 μM) were used to induce cardiac mesoderm from day 0 till day 3. XAV-939 (XAV, 5 μM) and VEGF (50 ng/ml) were used to induce CMECs and early cardiomyocytes (CMs) from day 3 to day 6. For paraxial mesoderm-ECs (PMECs), we adapted a protocol developed by Loh et al. (Loh et al., 2016)(Figure 1B). Briefly, high CHIR (8 μM) was used for the first two days followed by XAV (5 μM) for one day to induce posterior presomitic mesoderm (pPSM) on day 3 and low CHIR (1.5 μM) was used to induce anterior presomitic mesoderm (aPSM) from day 3 to day 5. VEGF was added to induce PMECs from day 5 to day 6. In order to characterize paraxial mesoderm differentiation and the PMEC lineage we established a double fluorescent hiPSC reporter line (NCRM1 PAX3^Venus^MSGN1^mCherry^) (Figure S1A). CRISPR/Cas9 assisted gene editing was used to target the fluorescent protein Venus to the PAX3 locus leading to transcriptional control by the endogenous PAX3 regulatory elements. The MSGN1^mCherry^ reporter was generated using a BAC construct integrated into the cells using the piggyBac transposon system. Flow cytometry analysis at different stages of paraxial mesoderm differentiation showed efficient induction of pPSM (MSGN-mCherry positive cells) and aPSM (PAX3-Venus positive cells) on day 2-3 and day 5, respectively (Figure S1B). Using our paraxial mesoderm protocol, more than 70% of cells acquired MSGN1-mCherry expression on day 2-3 (Figure S1C) and more than 50% of cells acquired PAX3-Venus on day 6 (Figure S1D). We next confirmed induction of endogenous PAX3 protein expression by immunostaining with a PAX3-specific antibody (Figure S1E). Notably, more than 90% of cells were positive for PAX3 on day 5; this could be because of relatively weak endogenous expression of Venus that could not be detected in PAX3^low^ cells.

**Figure 1.**
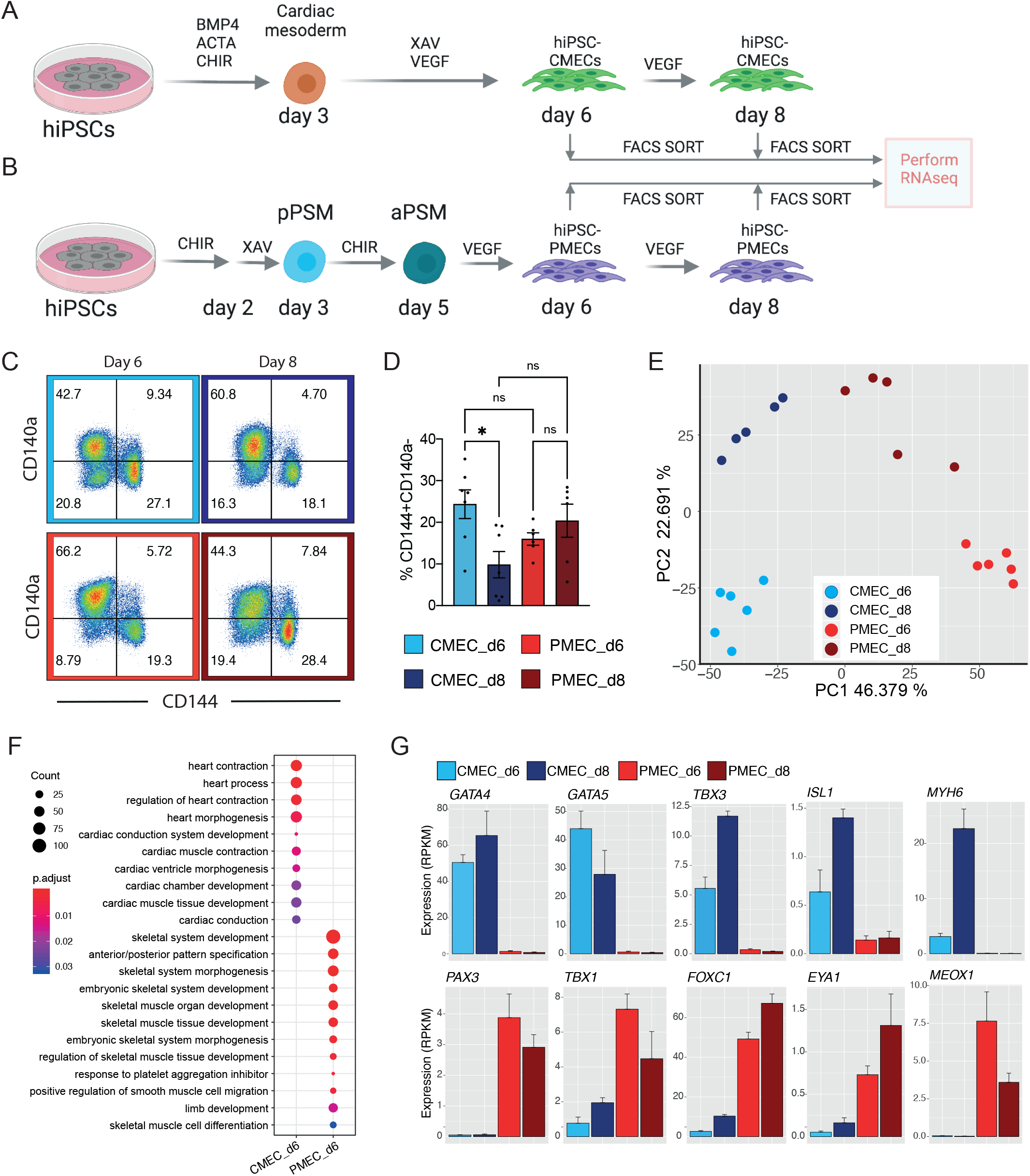
Characterization of ECs differentiated from hiPSCs using CMEC and PMEC protocols. (**A-B**) Schematic overview of CMEC (**A**) and PMEC (**B**) differentiation protocols. Endothelial cell (CD144+CD140a-) were FACS sorted on days 6 and day 8 for bulk RNA-seq. ACTA: activin-A. CHIR: CHIR99021. pPSM/aPSM: posterior/anterior presomitic mesoderm. LPM: lateral plate mesoderm. (**C**) Flow cytometry analysis of CD140a and CD144 expression on day 6 and day 8 of CMEC and PMEC differentiation. (**D**) Quantification of VEC (CD144)+CD140a-cells on day 6 and day 8 of CMEC and PMEC differentiation. Error bars represented standard deviations calculated from five to six independent differentiations. (**E**) PCA analysis of ECs sorted on day 6 and 8 of the CMEC or PMEC protocol. (**F**) GO term enrichment analysis for DEGs between CMECs and PMECs on day 6 of differentiation. Complete list of GO terms can be found in Table S2. Color represents the padjusted of enrichment analysis and dot size represents the count of genes mapped to the GO term. (**G**) Normalized gene expression levels (RPKM) of cardiac and skeletal related genes in CMECs and PMECs on day 6 and 8.

Gene expression analysis further confirmed comparable expression of pan-mesoderm markers *TBXT* and *MIXL1* in both cardiac and paraxial mesoderm differentiation conditions. On the other hand, expression of cardiac genes (*MESP1, GATA4* and *NKX2-5*) was restricted to cardiac mesoderm differentiation conditions and expression of paraxial mesoderm genes (*MSGN1, TBX6, PAX3*) was restricted to paraxial mesoderm differentiation conditions (Figure S1F).

Both cardiac and paraxial mesoderm differentiation conditions resulted in comparable percentages of CD144+CD140a-ECs on day 6 and day 8 of differentiation (Figure 1C-D). CD144+CD140a-ECs were sorted on day 6 and day 8 of differentiation from both cardiac and paraxial mesoderm conditions and underwent RNA sequencing (RNA-seq). Principle component analysis (PCA) showed that CMECs and PMECs clustered separately along PC1, and day 6 and day 8 were separated along PC2 (Figure 1E). On day 6, 3307 and 2592 genes were significantly differentially upregulated (FDR<0.05, fold-change>2) in CMECs and PMECs respectively (Table S1). Gene ontology (GO) analysis showed that cardiac related genes were specifically upregulated in day 6 CMECs (CMEC_D6), while genes related to skeletal system development and function were specifically upregulated in day 6 PMECs (PMEC_D6) (Figure 1F, Table S2). Genes involved in heart development, like *GATA4, GATA5, TBX3, ISL1* and *MYH6*, were highly expressed in day 6 and day 8 CMECs. *TBX3, ISL1* and *MYH6* were upregulated from day 6 to day 8 in CMECs (Figure 1G). Essential genes for skeletal muscle development like *PAX3, TBX1, FOXC1, EYA1* and *MEOX1* were largely expressed in day 6 and day 8 PMECs. *FOXC1* and *EYA1* were upregulated from day 6 to day 8, while *TBX1* and *MEOX1* were downregulated (Figure 1G). In summary, unbiased expression analysis by bulk RNA-seq showed differential gene expression signatures of cardiac and paraxial mesoderm derived ECs that corresponded to their known expression profiles *in vivo*.

### Reconstruction of the differentiation trajectories of ECs by single-cell RNA-seq

Having demonstrated that the two differentiation protocols result in ECs with distinct characteristics, we undertook an unbiased analysis of the complete cell population and reconstructed the EC differentiation trajectories. To this end, we performed scRNA-seq on day 6 of CMEC and PMEC differentiation from two independent biological replicates (Figure 2A-B, Figure S2A-B). The replicates appeared highly similar in a low-dimensional representation (Figure S2C) and were therefore combined for further analysis. Any remaining, undifferentiated hiPSCs were excluded from further analysis (Figure S2D-E). In the CMEC differentiation data set, cells were grouped into 3 clusters (cardiac mesoderm, cardiomyocytes and CMECs), as established previously (Cao et al., 2022) (Figure 2C). The three cell types were identified by known marker genes (Figure 2D, S3A-B, Table S3). The cardiac mesoderm cluster was characterized by mesoderm and early cardiac genes, such as *MESP1, SMARCD3, ABLIM1, TMEM88, ISL1, MYL5*, as well as the cell cycle-related genes *CDK6* and *NEK2*. The CMEC cluster was characterized by EC markers (*CDH5, CD34, KDR, HEY2, TEK, TIE1, ACVRL1, SOX17, ENG, ICAM2, PECAM1*).Cardiomyocytes were identified by expression of cardiomyocyte-associated genes, including *MYL4, TNNI1, MYL7, ACTA2, TNNT2, HAND2* and *NKX2-5*. To reveal the differentiation trajectories of the cells, we calculated the diffusion pseudotime using a cardiac cell mesoderm cell as root (Figure 2E). Pseudotime increased towards CMECs and cardiomyocytes, suggesting that both cell types differentiated from a common cardiac mesoderm progenitor.

**Figure 2.**
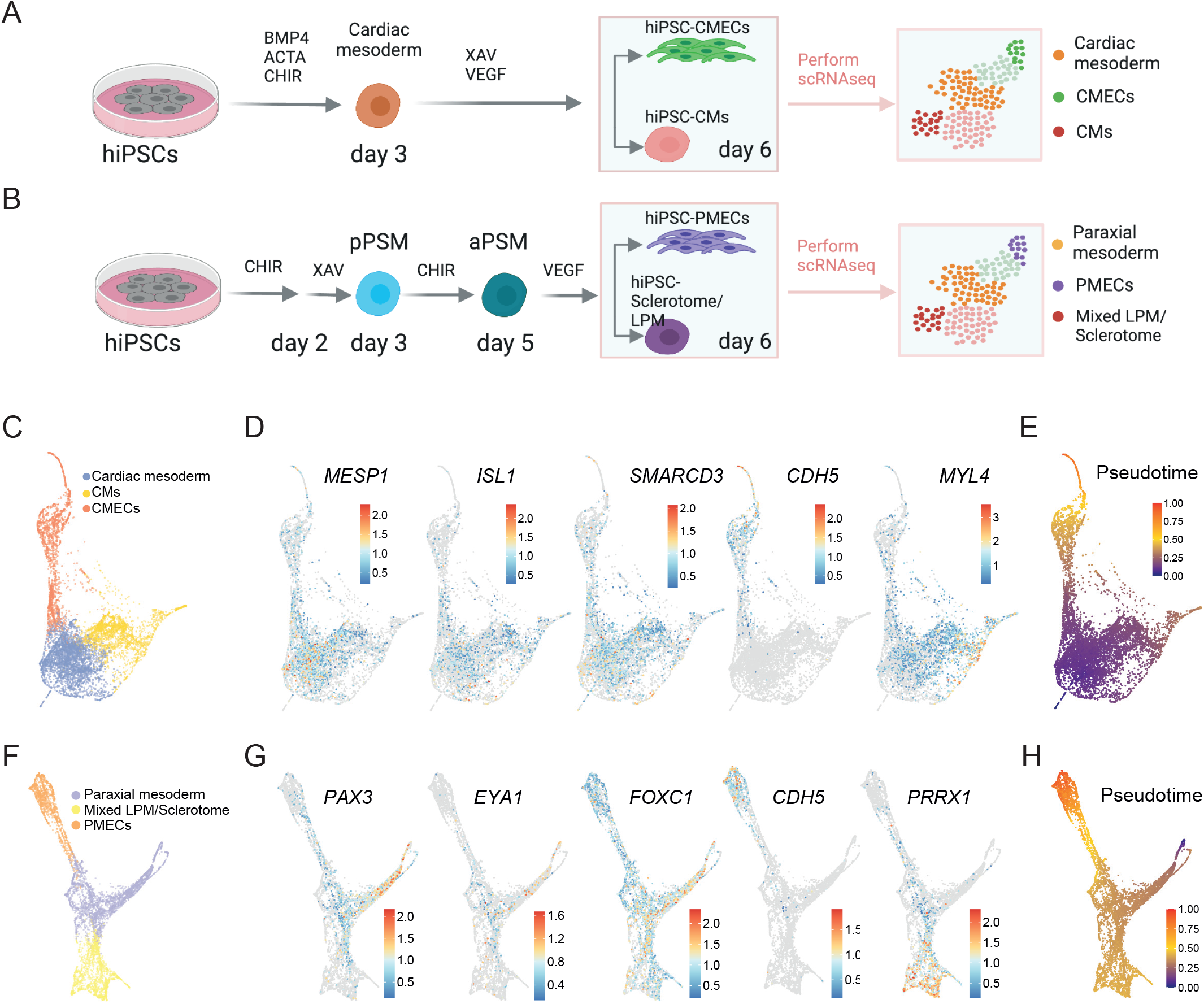
Single-cell RNA sequencing analysis of ECs differentiated from cardiac and paraxial mesoderm. (**A-B**) Schematic overview of CMEC (**A**) and PMEC (**B**) differentiation protocols until day 6. Cells were collected for scRNA-seq on day 6. ACTA: activin-A. CHIR: CHIR99021. pPSM/aPSM: posterior/anterior presomitic mesoderm. LPM: lateral plate mesoderm. (**C**) scRNA-seq of CMEC differentiation on day 6 (PAGA plot). Three clusters of cells, indicated by color, were identified. (**D**) PAGA plots show expression of *MESP1, ISL1, SMARCD3, CDH5, MYL4* on day 6 of CMEC differentiation. Color represents log transformed expression. (**E**) Diffusion pseudotime analysis of CMEC differentiation on day 6. (**F**) scRNA-seq of PMEC differentiation on day 6 (PAGA plot). Three clusters of cells, indicated by color, were identified. (**G**) PAGA plots show expression *PAX3, EYA1, FOXC1, CDH5, PPRX1* on day 6 of CMEC differentiation. Color represents log transformed expression. (**H**) Diffusion pseudotime analysis of PMEC differentiation on day 6.

In the PMEC differentiation data set, all cells were divided into 3 clusters (Figure 2F), which were interpreted as being paraxial mesoderm, PMECs and mixed lateral plate mesoderm (LPM)/sclerotome using marker gene analysis (Figure S3C, D, Table S4). The paraxial mesoderm cluster was characterized by expression of aPSM and dermomyotome genes, such as *MEOX1, PDGFRB, SIX1, CRABP2, NR2F1, EYA1, FOXC1 and PAX3*. PMECs were characterized by EC markers, like *ETV2, CDH5, CD34, KDR, ENG, SOX17, PLVAP, APLN, NRP1*. The mixed LPM/sclerotome cluster was characterized by LPM and sclerotome specific genes, such as *TMEM88, HAND1, TNNI1, PRRX1, ACTA2, DES, FOXH1, LEF1* and *JAG1* (Figure 2G, S3C-D). Diffusion pseudotime rooted in the paraxial mesoderm increased towards both PMECs and LPM/Sclerotome (Figure 2H). Both cell types therefore likely differentiated from a common paraxial mesoderm progenitor.

### Acquisition of an organ-specific identity in cardiac microtissues

Being able to produce ECs with properties corresponding to their mesodermal origins enabled us to test in how far the cellular microenvironment can either reinforce or reverse this specification i.e. how “plastic” the ECs are. Specifically, we set out to mimic the cardiac microenvironment *in vitro* using a protocol for creating cardiac MTs, published previously by our group (Giacomelli et al., 2020). Briefly, CD34+ CMECs or PMECs were sorted on day 6 and combined with hiPSC-derived cardiomyocytes (hiPSC-CMs) and hiPSC-derived fibroblasts (hiPSC-CFs) in a ratio of 15:70:15 to form MTs. MTs made from CMECs (CM_MTs) and PMECs (PM_MTs) were collected after 21 days from two independent biological replicates by scRNA-seq (Figure 3A, Figure S2A-B). The replicates appeared highly similar in a low-dimensional representation (Figure S2C) and were therefore combined for further analysis. Any remaining, undifferentiated hiPSCs were excluded from further analysis (Figure S2F-G). Both CM_MTs and PM_MTs datasets were divided into three clusters that correspond to hiPSC-CFs, hiPSC-CMs and hiPSC-ECs (Figure 3B). Marker genes identified for each cluster, confirmed the cluster identities (Table S6).

**Figure 3.**
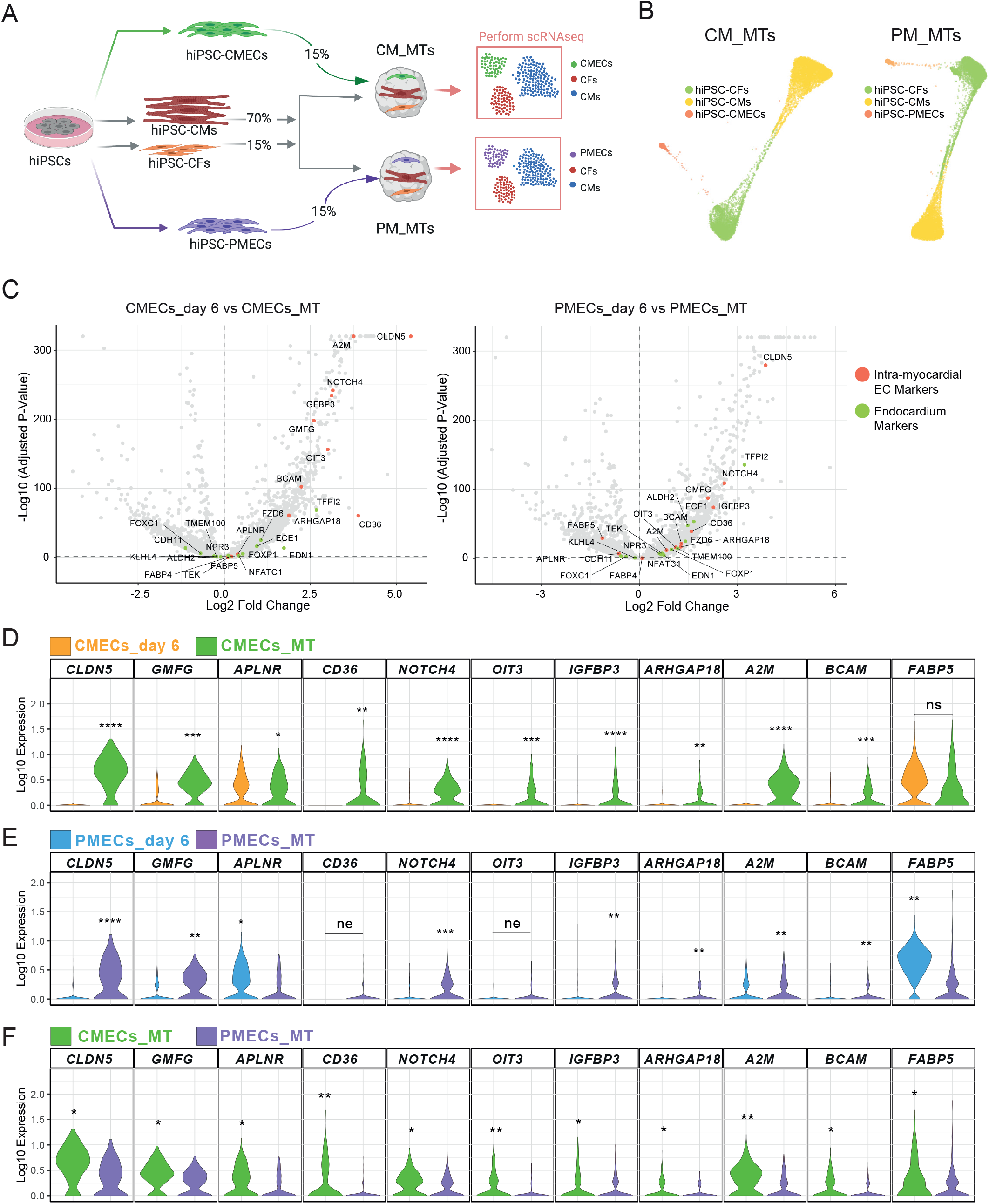
hiPSC-ECs acquired organ-specific signatures in a cardiac microenvironment. (**A**) Generation of cardiac MTs from hiPSC-CMs, hiPSC-CFs and hiPSC-ECs. CMECs and PMECs were used for CM_MTs and PM_MTs respectively. MTs were collected after 21 days for scRNAseq. (**B**) scRNAseq data of CM_MTs (left) and PM_MTs (right) were visualized using PAGA. Three clusters of cells were identified in both datasets. (**C**) Volcano plot showing fold changes and p-values of differential expression tests between CMECs_day 6 and CMECs_MTs (left), or PMECs_day 6 and PMECs_MTs (right). Representative intra-myocardial and endocardial markers that are differentially expressed (p_adjusted_<0.05) are highlighted in red and green, respectively. (**D-F**) Violin plots of gene expression in PMECs_ day 6, CMECs_day6, CMECs_MT and PMECs_MT for representative Intra-myocardial EC markers. Asterisks indicate significance level of differential gene expression tests between the respective populations. ns: p>0.05; * p <= 0.05; ** p<= 1e-10; *** p<= 1e-100; **** p<= 1e-200. The clusters with higher expression value were labeled. ne: not expressed (0 counts) in >85% of cells in both groups.

In order to assess to what extent ECs in MTs acquired an organ-specific identity, we compared their expression profiles to primary ECs in a published data set of the human fetal heart (Asp et al., 2019) (Figure S4A). In this data set, we reannotated the original endothelium/pericytes/adventia cluster (cluster 10) as intramyocardial ECs, based on differentially expressed markers such as *A2M, CD36, APLNR, ARHGAP18, IGFBP3, CLDN5, FABP4* and *FABP5* (Figure S4B, Table S5). The cluster annotated as capillary endothelium (cluster 0) in the original publication was reannotated as endocardium, due to the presence of differentially expressed markers like *NPR3, ALDH2, CDH11, ECE1, TMEM100, FOXC1* and *EDN1* (Figure S4B, Table S5). Supporting the differential expression test, UMAP visualization of representative intra-myocardial and endocardial markers showed specific expression in the respective clusters (Figure S4C-D).

We next compared CMECs and PMECs on day 6 of differentiation (CMECs_day 6 and PMECs_day 6) with the CMECs and PMECs in MTs (CMECs_MT and PMECs_MTs) respectively. CMECs in MTs upregulated expression of intramyocardial makers, such as *CLDN5, GMFG, APLNR, CD36, NOTCH4, OIT3, IGFBP3, ARHGAP18, A2M* and *BCAM*, but not *FABP5*, compared to CMECs isolated on day 6 of differentiation (Figure 3C,D, Table S7). PMECs in MTs upregulated expression of a few intramyocardial markers, such as *CLDN5, GMFG, NOTCH4, IGFBP3, ARHGAP18, A2M* and *BCAM*, but not *APLNR, CD36, OIT3* and *FABP5* compared to PMECs isolated on day 6 of differentiation (Figure 3C,E, Table S7). We also found some endocardial markers upregulated in both CMECs and PMECs in MTs (*TFPI2, EDN1, ECE1, FOXP1)(S5A-*B, Table S7). However, the differences in endocardial marker expression were smaller compared to intramyocardial markers. Notably, the expression of several intramyocardial markers, especially *APLNR, CD36, OIT3, ARHGAP18, A2M, BCAM* and *FABP5* was higher in CMECs in MTs compared to PMECs in MTs (Figure 3F, Table S8). Although the expression of endocardial markers was also higher in CMECs in MTs compared to PMECs in MTs, their average expression levels were lower in general when compared to intramyocardial markers (Figure S5C, Table S8). Importantly, endocardial makers, including *CDH11, FOXC1, FZD6, TMEM100* and *NPR3*, were barely expressed in either CMECs or PMECs in MTs (Figure S5C). Overall, intramyocardial EC identify was acquired by all hiPSC-ECs in the cardiac MT environment.

### Distinct cell identities are preserved in cardiac microtissues composed of either cardiac or paraxial mesoderm-derived ECs

To obtain a clearer view of the similarities between ECs in MTs and primary fetal heart ECs, we merged the CM_MT and PM_MT datasets with the human fetal heart dataset (Asp et al., 2019) (Figure 4A-B). In the case of both the CM_MT and the PM_MT dataset, we found that CFs in MTs (CF_MT) clustered together with fetal heart fibroblast-like cells and CMs in MTs (CM_MT) clustered together with fetal heart ventricular CMs. Notably, both CMECs in MTs (CMECs_MT) and PMECs in MTs (PMECs_MT) clustered together with fetal heart intramyocardial ECs and not endocardium (Figure 4A-D). To quantify our observation, we calculated the distances (in expression space) between each cell in MTs and the fetal heart dataset. This calculation showed that CF_MT cells are closest to fibroblast-like cells *in vivo* (related to cardiac skeleton connective tissue), CM_MT cells are closest to ventricular cardiomyocytes and CMECs_MT as well as PMECs_MT are closest to intramyocardial ECs in human fetal heart (Figure 4E). Annotating the *in vitro* cells based on the closest *in vivo* neighbors revealed that cell type identities were very similar in CM_MTs and PM_MTs (Figure 4E).

**Figure 4.**
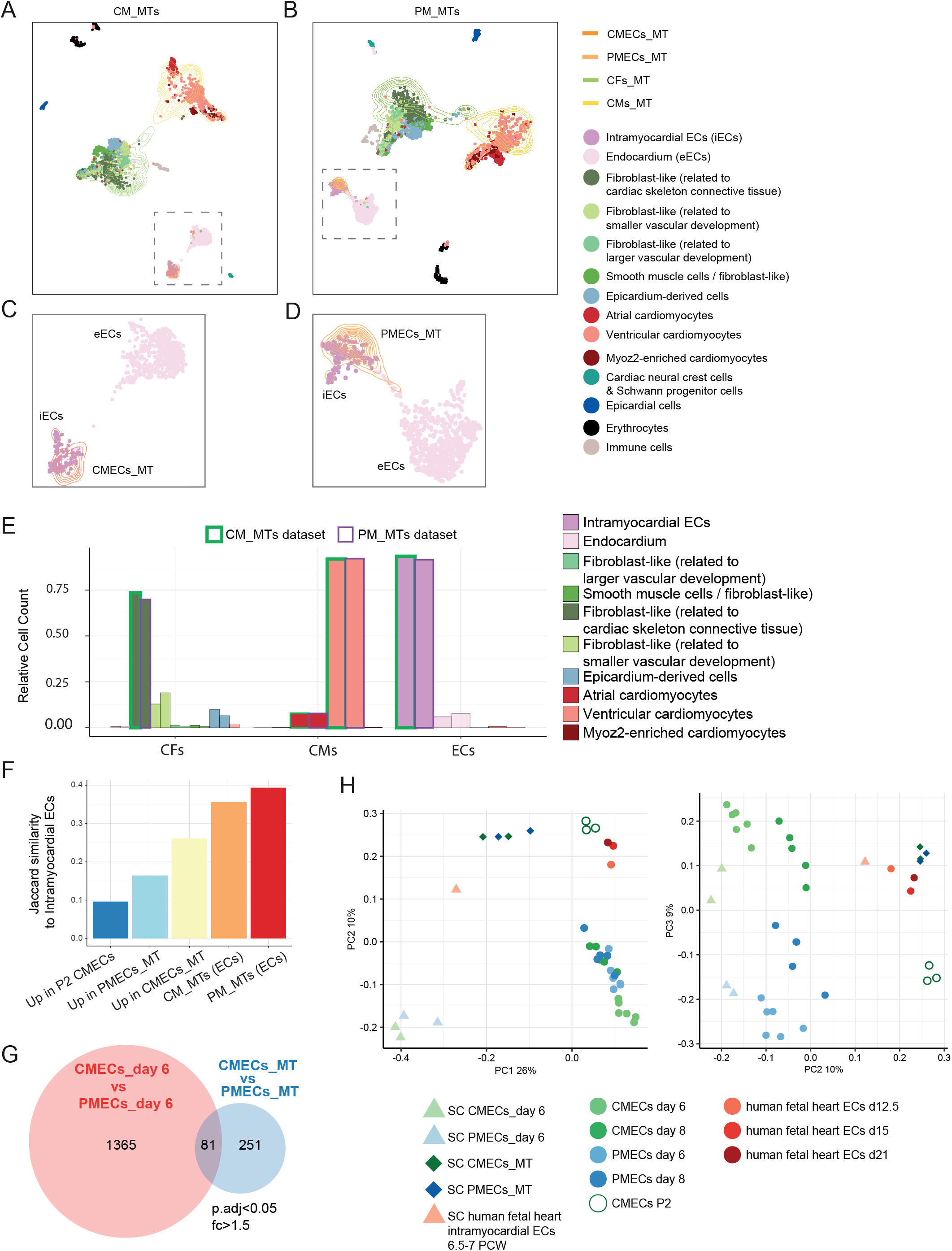
CMECs and PMECs acquired intramyocardial identity in cardiac microenvironment. (**A-B**) Low-dimensional representation (UMAP) of the human fetal heart data set (Asp et al., 2019) integrated with the CM_MTs (**A, C**) or PM_MTs (**B, D**) dataset. EC clusters are marked with squares. Cell clusters are indicated by color. Cells in fetal heart dataset are presented with dots and cells in MTs datasets are presented with contour lines. (**C-D**) Zoom-ins of the EC clusters marked in (**A-B**). (**E**) Annotation of CM_MT and PM_MT cells based on nearest neighbors in the *in vivo* dataset. Cells from CM_MTs and PM_MTs are outlined with a green or violet contour, respectively. (**F**) Jaccard similarity to intramyocardial ECs from the human fetal heart dataset was calculated for each indicated group of genes. CM_MTs (ECs): specific markers of cluster CMECs within the CM_MT dataset; PM_MTs (ECs): specific markers of cluster PMECs within the PM_MT dataset; Up in CMECs_MT: DEGs that are higher in CMECs_MT compared to CMECs_day 6; Up in PMECs_MT: DEGs that are higher in PMECs_MT compared to PMECs_day 6; Up in P2 CMECs: DEGs that are higher in passage two CMECs compared to CMECs_day 6. (**G**) Venn diagram shows numbers and overlap of DEGs (p_adjusted_<0.05 and fold-change > 1.5) between CMECs and PMECs from day 6 (in red) and MTs (in blue). (**H**) PCA plot of different EC populations in scRNAseq (triangle and diamond) and bulk RNA-seq (circle) datasets. Average expression values of all cells in the cluster were used for the scRNA-seq data.

Correspondingly, the set of markers of the EC cluster in CM_MTs and the PM_MTs, CM_MTs (ECs) and PM_MT (ECs) respectively, showed a high overlap (Jaccard similarity) with the markers of intra-myocardial ECs we extracted from the *in vivo* data set (Figure 4F). The gene set upregulated in CMECs_MT compared to CMECs_day 6 (Up in CMECs_MTs) had a higher overlap with intramyocardial EC markers than the set of genes upregulated in PMECs_MTs compared to PMECs_day 6 (Up in PMECs_MTs) (Figure 4F). For comparison, we also profiled CMECs that were cultured for two additional passages in monoculture. Genes that were upregulated in these cells compared to CMECs_day 6 (Up in P2 CMECs) overlapped the least with intra-myocardial EC markers (Figure 4F). This result excludes the possibility that the effects observed in MTs are simply due to environment-independent differentiation progression over time. CMECs_MT and PMECs_MT thus both resembled intramyocardial ECs but a difference between the two differentiation methods remained. To quantify this difference directly, we used differential gene expression analysis. On day 6 of differentiation, 1446 genes were differentially expressed between CMECs and PMECs (Table S9), while only 332 genes were differentially expressed between CMECs_MT and PMECs_MT (Table S8). 81 genes were shared between the two sets and 251 genes were differentially expressed only between CMECs_MT and PMECs_MT (Figure 4G). Intramyocardial marker genes (*CD36*, *OIT3, A2M*, *CLDN5*, *APLNR*, *FABP5*) were among these 251 genes that were differentially expressed between the CMECs_MT and PMECs_MT. This is also in line with our observations that expression of some intramyocardial marker genes were higher in CMECs in MTs compared to PMECs in MTs. Next, all EC clusters from bulk and single cell RNA-seq datasets were combined and visualized using principal component analysis (PCA) (Figure 4H). CMECs_day 6 and PMECs_day 6 clustered far apart, while CMECs_MT and PMECs_MT clustered closely together. Bulk and single cell RNA-seq samples clustered together for both CMECs and PMECs. CMECs_MT and PMECs_MT were found closer to fetal heart intramyocardial ECs and fetal heart ECs (Figure 4H) sequenced in our previous study (Giacomelli et al., 2020). Altogether, analysis of the *in vitro* and *in vivo* data sets demonstrated that the cardiac tissue microenvironment resulted in hiPSC-ECs acquiring an intramyocardial EC identity independent of their developmental origin and, further, gene expression differences due to distinct mesodermal origins were partially removed.

## DISCUSSION

In the present study we derived ECs from hiPSCs from two mesoderm lineages namely LPM and PM, as previous lineage tracing studies have shown that LPM and PM serve as a major source of ECs in the developing embryo (Lagha et al., 2009; Mayeuf-Louchart et al., 2016; Pardanaud et al., 1996). We showed that ECs isolated on day 6 and day 8 of differentiation retained their developmental lineage history. Transcription factors involved in the heart and skeletal muscle development were highly expressed in CMECs and PMECs, respectively. Additional approaches, including scRNA-seq, showed lineage diversification from a common cardiac and paraxial mesoderm progenitor during differentiation of CMECs and PMECs, respectively. At the same time, an organ-specific EC signature was absent in ECs differentiated either from cardiac or paraxial mesoderm on day 6 and day 8 of differentiation, or upon extended culture.

Local microenvironmental cues result in acquisition of organ-specific characteristics in ECs (Aird, 2012). To model the influence of cell-extrinsic factors, we took advantage of our cardiac MT model which mimics the heart-specific microenvironment, as it integrates CMs, CFs and ECs (Giacomelli et al., 2020). Although both CMECs and PMECs acquired an iEC identity after incorporation into MTs, several intramyocardial EC markers (*APLNR, CD36, OIT3, ARHGAP18, A2M, BCAM* and *FABP5*) were more strongly upregulated in CMECs compared to PMECs in cardiac MTs. A recent study in mouse embryos showed that transcriptional heterogeneity in the sinus venosus (SV) and endocardium-derived iECs declines over time (Phansalkar et al., 2021). Therefore, it would be interesting to investigate whether PMECs require longer culture in cardiac MTs to acquire comparable expression of intramyocardial EC markers as in CMECs. On the other hand, endocardial markers were not detected in either CMECs or PMECs in cardiac MTs. This is in line with previous evidence that the cardiac MTs microenvironment recapitulates myocardial and not endocardial layers of the heart. Importantly, CFs and CMs in both CM_MT and PM_MT datasets were highly similar and clustered together with fetal heart fibroblast-like cells and fetal heart ventricular CMs respectively. This showed that developmental origin of ECs does not influence CF or CM identities in cardiac MTs.

Among EC lineages, PM serves a source of lymphatic ECs in the heart, skin and lymph node (Lenti et al., 2022; Lupu et al., 2022; Stone and Stainier, 2019). PMECs showed increased expression of genes important for the development of lymphatic vasculature, such as *TBX1, FOXC1, LYVE1, VEGFC*. At the same time, expression of the master regulator of lymphatic EC differentiation (*PROX1*) was not detected in PMECs either at day 6 or in cardiac MTs. It would be interesting to explore whether addition of known lymphatic EC growth factors promotes *PROX1* expression in PMECs.

Genetic lineage tracing in mice has identified a variety of developmental origins for organ-specific ECs in different tissues. However, whether developmental origin is a prerequisite-or simply a default developmental route remains an open question. On the other hand, the present and previous studies demonstrate that local microenvironmental cues might play a bigger role in not only the acquisition but also the maintenance of the organ-specific characteristics. This is also in line with recent study on sinusoidal ECs of the liver (Gómez-Salinero et al., 2022). Earlier studies have demonstrated the importance of the *Gata4* transcription factor in the differentiation of liver ECs and the acquisition of sinusoidal-like identity (Géraud et al., 2017; Zhang et al., 2016b). However, scRNA-seq analysis of liver ECs showed that *Gata4* is not restricted to sinusoidal ECs but it is expressed by all ECs in the liver (Gómez-Salinero et al., 2022). Instead, the *c-Maf* transcription factor was restricted to sinusoidal liver ECs and was regulated by BMP9 that is highly expressed by hepatic stellate cells (HSC) (Breitkopf-Heinlein et al., 2017). Furthermore, overexpression of c-Maf was sufficient to induce sinusoidal-like characteristics in human umbilical vein ECs (HUVECs). The same is true for heart ECs that are derived from multiple developmental pools of EC progenitors that converge to a similar state over time (Milgrom-Hoffman et al., 2011; Phansalkar et al., 2021; Sharma et al., 2017b).

In summary, we demonstrated that ECs, derived from two distinct mesoderm lineages namely LPM and PM, acquire organ-specific characteristics upon incorporation into the cardiac MT environment which includes CMs and CFs. We expect that our findings will guide the derivation of organ-specific ECs from hiPSCs in the future and lay the foundation for various biomedical applications, from creating of disease models to transplantation therapies.

## Materials and Methods

### hiPSC culture

hiPSC lines LUMC0020iCTRL-06 and NCRM1 (NIH Center for Regenerative Medicine NIH CRM, obtained from RUDCR Infinite Biologics at Rutgers University) were cultured in in TeSR-E8 on Vitronectin XF and were routinely passaged once a week using Gentle Cell Dissociation Reagent (all from Stem Cell Technologies, Vancouver, Canada).

### Generation of PAX3^Venus^MSGN1^mCherry^ hiPSC dual reporter line

Prior to targeting, NCRM1 hiPSCs were passaged as a bulk on feeders in hESC-medium (Costa et al., 2007). RevitaCell (Life Technologies, Carlsbad, CA, USA) was added to the medium (1:200) after every passage to enhance viability after single cell passaging with TrypLE (Life technologies). PAX3^Venus^ was generated by CRISPR/Cas9 as follow: NCRM1 hiPSCs were passaged with ratio 1:3 into 60 mm dishes to reach 60-70% confluency the next day for transfection. Cells were transfected with pCas9-GFP (Addgene plasmid #44719), pBR322-U6-hPAX3-gRNA-S1 containing sgRNA CCGGCCAGCGTGGTCATCCT and repair template p15A-cm-hPAX3-Venus-neo-1kb containing a Venus-neo cassette with 1 kb hPAX3 homology arms. The antibiotic selection marker is flanked by FRT sites for Flp-mediated excision. 20 μl lipofectamine (Invitrogen, Waltham, Massachusetts, USA), 8 μg of pCas9-GFP, 8 μg of sgRNA plasmid and 8 μg of linearized repair template were diluted in 600 μl of Opti-MEM and added to each 60 mm dish. After 18 hours the medium was changed to hESC medium. After another 6 hours G-418 (50 μg/ml) selection was started and was kept for 1 week. Surviving cells were cultured in hESC medium, passaged and transferred into 6-well plates for the transfection of Flp recombinase expression vector to remove the neomycin cassette. 300 μl of Opti-MEM containing 10 μl lipofectamine and 4 μg CAGGs-Flpo-IRES-puro plasmid was added per well for 18 hours. Puromycin (0,5 μg/ml) selection was started 24 hours post transfection and lasted for 2 days. Once recovered, cells were passage into 96-well format for clonal expansion via limited dilution. Targeted clones were identified by PCR and Sanger sequencing (BaseClear, Leiden, Netherlands). The MSGN1^mCherry^ reporter line was generated by Transposon mediated BAC transgenesis using protocols described by (Rostovskaya et al., 2012). In brief, a human BAC (RP11-12L16) with piggyBac transposon repeats flanking the bacterial backbone and with mCherry inserted directly after the initiating Methionine of MSGN1 was transfected together with a piggyBac Transposase into NCRM1 hiPSCs.

### Differentiation of ECs from cardiac and paraxial mesoderm

ECs from cardiac mesoderm were differentiated as previously described (Giacomelli et al., 2017). ECs from paraxial mesoderm were differentiated using a modified Loh et al. protocol (Loh et al., 2016). Briefly, 5 x 10^4^ cells per cm^2^ were seeded on plates coated with 75 μg/mL Matrigel (growth factor reduced) (Corning) the day before differentiation (day −1). At day 0, paraxial mesoderm was induced by changing TeSR-E8 to BPEL (Bovine Serum Albumin [BSA], Polyvinyl alcohol, Essential Lipids) medium (Ng et al., 2008), supplemented with 8 μM CHIR99021. At day 2, cells were refreshed with BPEL supplemented with 5 μM XAV939. At day 3, cells were refreshed with BPEL supplemented with 4 μM CHIR99021. From day 5 onwards, cells were refreshed every 3 days with BPEL medium supplemented with 50 ng/ml VEGF.

### Fluorescence-activated cell sorting

Cells were dissociated with TrypLE on day 6 and 8 of CMEC and PMEC protocol and stained with VE-Cadherin PE-conjugated Antibody (R&D Systems). Then VEC+ cells were sorted using FACSAria III. Total RNA was extracted right after sorting using the NucleoSpin^®^ RNA kit (Macherey-Nagel, Düren, Germany).

### Generation of 3D Cardiac Microtissues (MTs)

Cardiac MTs were generated from hiPSC-derived ECs, CFs and CMs as previously described (Giacomelli et al., 2020). Briefly, on day 6 of CMEC and PMEC differentiation, CD34^+^ ECs were isolated using a Human cord blood CD34 Positive selection kit II (StemCell Technologies) following the manufacturer’s instructions. On the day of MT formation, freshly isolated hiPSC-ECs and cultured hiPSCs-CFs and hiPSC-CMs were combined together to 5000 cells (70% cardiomyocytes, 15% endothelial cells and 15% cardiac fibroblasts) per 50μl BPEL medium supplemented with VEGF (50 ng/ml) and FGF2 (5 ng/ml). Cell suspensions were seeded on V-bottom 96 well microplates (Greiner bio-one, Kremsmünster, Austria) and centrifuged for 10 min at 1100 rpm. MTs were incubated at 37°C, 5% CO_2_ for 21 days with media refreshed every 3-4 days. scRNAseq analysis of MTs was performed after 21 days.

### Fluorescence-activated cell sorting

Cells were detached using TrypLE for 5 min at 37°C and washed once with FACS buffer (PBS containing 0.5% BSA and 2 mM EDTA). Primary antibodies CD144 (1:50, eBioscience), CD140a (1:20, BD Bioscience) were added for 1 hr at 4°C. Samples were measured on MACSQuant VYB (Miltenyi Biotech) equipped with a violet (405 nm), blue (488 nm) and yellow (561 nm) laser. The results were analyzed using Flowjo v10 (FlowJo, LLC). For FACS sorting on day 6 and day 8 of the CMEC and PMEC protocol, CD144+CD140a-cells were sorted using FACSAria III (BD-Biosciences).

### Quantitative Real-Time Polymerase Chain Reaction (qPCR)

Total RNA was extracted using the NucleoSpin^®^ RNA kit according to the manufacturer’s protocol. cDNA was synthesized using an iScript-cDNA Synthesis kit (Bio-Rad, Hercules, CA, USA). iTaq Universal SYBR Green Supermixes (Bio-Rad) and Bio-Rad CFX384 real-time system were used for the PCR reaction and detection. Relative gene expression was determined according to the standard ΔCT calculation and normalized to the housekeeping gene RPL37A.

### Bulk RNA sequencing and analysis

Bulk RNAseq of passage two CMECs (CMECs P2) and human fetal heart ECs at gestation age Week(W)12.5, W15 and W21 were performed in our previous study (Giacomelli et al., 2020) and obtained from GEO accession number GSE116464.

Bulk RNAseq of day 6 and 8 of CMEC and PMEC differentiation were performed at BGI (Shenzhen, China) using the Illumina Hiseq4000 (100bp paired end reads). Raw data was processed using the LUMC BIOPET Gentrap pipeline (https://github.com/biopet/biopet), which comprises FASTQ preprocessing, alignment and read quantification. Sickle (v1.2) was used to trim low-quality read ends (https://github.com/najoshi/sickle). Cutadapt (v1.1) was used for adapter clipping (Martin, 2011), reads were aligned to the human reference genome GRCh38 using GSNAP (gmap-2014-12-23) (Wu and Nacu, 2010; Wu and Watanabe, 2005) and gene read quantification with htseq-count (v0.6.1p1) against the Ensembl v87 annotation (Yates et al., 2015). Gene length and GC content bias were normalized using the R package cqn (v1.28.1) (Hansen et al., 2012).Genes were excluded if the number of reads was below 5 reads in ≥90% of the samples.

Differentially expressed genes were identified using generalized linear models as implemented in edgeR (3.24.3) (Robinson et al., 2009). P-values were adjusted using the Benjamini-Hochberg procedure and FDR ≤ 0.05 was considered significant. Analyses were performed using R (version 3.5.2). PCA plot was generated with the built-in R functions prcomp using transposed normalized RPKM matrix. Correlation among samples was calculated using cor function with spearman method and the correlation heatmap was generated with aheatmap function (NMF package). Gene ontology enrichment was performed using compareCluster function of clusterProfiler package (v3.10.1) (Yu et al., 2012) and q ≤ 0.05 was considered significant.

### Single-cell RNA sequencing and analysis

#### Library preparation and sequencing

Library preparation was performed as previously described (Giacomelli et al., 2020). Briefly, single cells were loaded into the 10X Chromium Controller for library construction using the Single-Cell 3’ Library Kit, Version 2 Chemistry (10x Genomics, Pleasanton, CA, USA) according to the manufacturer’s protocol. Next, indexed cDNA libraries were sequenced on the HiSeq4000 platform. Single-cell expression was quantified using unique molecular identifiers (UMIs) by 10x Genomics’ “Cell Ranger” software.

The mean reads per cell for all eight data sets: CMEC (R1): 28,499; CMEC (R2): 29,388; PMEC (R1): 31,860; PMEC (R2): 38,415; CM_MT (R1): 39,319; CM_MT (R2): 29,741; PM_MT (R1): 36,726; PM_MT (R2): 26,421.

#### Single-cell RNAseq data pruning and normalization

For data pruning and normalization, the two replicates of each of the 4 conditions (CMEC, PMEC, CM_MT and PM_MT) were combined without batch correction. Then, cells with a low number of genes per cell (1200 [CMEC], 1200 [PMEC], 900 [CM_MT], 750 [PM_MT], see Fig. S2A-B) were removed. Genes expressed in less than 2 of the remaining cells with a count of at most 1 were excluded from further analysis. Each combined data set was normalized using the R package scran (V 1.14.6) (Lun et al., 2016). Highly variable genes (HVGs) were calculated (improvedCV2 from the scran package) for each replicate of the combined data sets after excluding ribosomal genes [Ribosomal Protein Gene Database], stress markers (Brink et al., 2017) and mitochondrial genes. For downstream analysis the top 5% HVGs were used after excluding proliferation (Whitfield et al., 2006) and cell cycle (Giotti et al., 2017) related genes.

#### Cell cycle analysis and batch correction

For each combined data set, cell cycle analysis was performed with the scran package using the cyclone function (Scialdone et al., 2015) on normalized counts (Figure S2H). Cells with a G2/M score higher than 0.2 were considered to be in G2/M phase. Otherwise, they were classified as G1/S. Using this binary classifier as predictor, we regressed out cell cycle effects with the R package limma (V 3.42.2) (Ritchie et al., 2015) applied to log-transformed normalized counts. Then, for each combined data set, the two replicates were batch corrected with fast mutual nearest neighbors correction method (MNN) (Haghverdi et al., 2016) on the cell cycle corrected counts, using the 30 first principal components and 20 nearest-neighbors (Figure S2C).

#### Clustering

For each combined data set, batch-corrected counts were standardized per gene and then used to create a shared nearest neighbour (SNN) graph with the *scran* R package (d = 30, k =2). Louvain clustering was applied to the SNN graph using the *igraph* python package (V 0.7.1) with these resolution parameters: 0.4 [CMEC], 0.4 [CM_MT], 0.3 [PMEC], 0.1 [PM_MT]. For the CMEC data set, this resulted in 5 clusters (Figure S2D). Two of these 5 clusters were excluded from further analysis based on the expression of pluripotency markers (Figure S2E). For the PMEC data set, this resulted in 3 clusters (Figure 2F). For CM_MT and PM_MT, clustering resulted in 4 clusters (Figure S2F and S2G), where one cluster was excluded from further analysis, because it was mainly present in one of the two replicates. Additionally, the attempt to map this cluster to in vivo data, resulted in mostly unassigned cell types (plot not shown). For PMEC, clustering resulted in 3 clusters.

#### Dimensionality reduction and pseudotime

Dimensionality reduction was performed using the python *scanpy* pipeline (V 1.4.6). For both data sets, CMEC and PMEC, a 20 nearest-neighbors (knn, k=20) graph was created from diffusion components of the batch corrected data sets. Diffusion components are the eigenvectors of the diffusion operator which is calculated from Euclidean distances and a gaussian kernel. The aim is to find a lower dimensional embedding which considers the cellular progression. Both graphs were projected into two dimensions with the default force-directed graph layout and starting positions obtained from the partition-based graph abstraction (PAGA) output (Wolf et al., 2019). PAGA estimates connectivities between partitions and performs an improved version of diffusion pseudotime. Diffusion pseudotime (Haghverdi et al., 2016; Wolf et al., 2019) was calculated on these graphs with root cells selected based on the graph layout from the “Cardiac Mesoderm” cluster in CMEC, and the “Paraxial Mesoderm” cluster in PMEC.

For CM_MT and PM_MT, the knn graphs (k=50 for PM_MT, k=100 for CM_MT) were created from the first 30 principal components of the batch corrected data sets. These graphs were projected into two dimensions with the default force-directed graph layout and starting positions from the PAGA output.

#### *In vivo* data analysis and mapping

The *in vivo* data set, downloaded from https://www.spatialresearch.org/resources-published-datasets/doi-10-1016-j-cell-2019-11-025/, contains a 6.5 PCW human embryonic cardiac tissue sample. The clusters and cluster annotations were obtained from the original publication (Asp et al., 2019). The data set was normalized with the scran R package and HVGs were calculated as described in section “Single-cell RNA-seq data pruning and normalization”. Dimensionality reduction was performed with the R package *umap* (V 0.2.5.0) using 20 nearest-neighbors, min_dist = 0.7 and Euclidean distance.

#### Differential expression analysis

All differential expression tests were performed with edgeR (V 3.28.1) (Robinson et al., 2009) using a negative binomial regression and raw counts. The predictors in the regression were: cluster and replicate (both discrete variables), as well as the total number of counts per cell.

For marker gene analysis (Figures S3A and S3C), p-values were obtained for a contrast between the cluster of interest and all other clusters using regression coefficients averaged over the replicates. For tests between different data sets (Figure 3C), the corresponding endothelial cell cluster was extracted from each data set. Then, a contrast between MT and day 6 was calculated by averaging over the predictors of both replicates. For the *in vivo* test (Figure S4B), intra-myocardial EC and endocardium clusters were extracted from the data set to calculate the contrast between them. P-values were adjusted for multiple hypothesis testing with the Benjamini-Hochberg method.

#### Comparison to the *in vivo* data set

CM_MT and PM_MT data sets were mapped on the *in vivo* data set using the MNN method (d = 30 principal components, k = 100 nearest neighbors). First, *in vitro* replicates were mapped to each other, then the *in vivo* data was mapped on the combined *in vitro* data, using normalized, log-transformed counts and the 10% top HVGs of the *in vivo* data set. Dimensionality reduction was performed with the R package *umap* using 100 nearest-neighbors, min_dist = 0.3 and Euclidean distance. K-nearest-neighbour (KNN) assignment was performed in the batch corrected, principal component space (30 PCs). The 100 nearest-neighbors in the *in vivo* data set based on Euclidean distances were calculated for each *in vitro* cell. The *in vitro* cell was ascribed the cell type most abundant among the 100 *in vivo* neighbors. Each such assignment received a confidence score, which is the number of *in vivo* neighbors with that cell type divided by the number of all nearest neighbors (=100). A cell was not ascribed a cell type if either the average distance to its nearest neighbour exceeded a certain threshold (determined by the long tail of the histogram of average distances: 0.35), or the assignment had a confidence score less than 0.5. In addition, clusters containing less than 10 cells were not ascribed a cell type.

For the Jaccard similarity measure, marker genes of each differential expression test were selected with adjusted p-value < 0.05. The remaining genes were ranked by log2 fold change and the first 478 genes were selected for analysis. Then, the Jaccard distances were calculated between the marker genes of intra-myocardial endothelial cells and each of the other gene sets.

For principal component analysis (Figure 4H), human fetal bulk samples and in vitro bulk samples (Giacomelli et al., 2020) were combined with the single cell data sets. For each single-cell data set, the endothelial cells were extracted and the sum per gene over all cells was calculated. Then, bulk and single cell samples were log-transformed and combined into one data set. Principal component analysis was applied on the gene-wise standardized data set, using marker genes of the intra-myocardial endothelial cells from the *in vivo* data set.

## Data Availability Statement

The accession numbers for the bulk and single cell RNA sequencing datasets reported in this paper are https://www.ncbi.nlm.nih.gov/geo/ GEO: GSE151427 (day 6 and day 8 CMEC and PMEC (bulk); day 6 PMEC (single cell); PM_MT (single cell)); GSE202901 (day 6 CMEC (single cell)); GSE147694 (CM_MT (single cell)).

## Statistics

Statistical analysis was conducted with GraphPad Prism 7 software (San Diego, CA, USA). Data are represented as mean± SD.

## Acknowledgements

This project received funding from the European Union’s Horizon 2020 Framework Programme (668724); Netherlands Organ-on-Chip Initiative, an NWO Gravitation project funded by the Ministry of Education, Culture and Science of the government of the Netherlands (024.003.001); the European Union’s Horizon 2020 and innovation program under a Marie Sklodowska-Curie grant agreement no. 707404. We thank Elisa Giacomelli and Milena Bellin for generation of cardiac microtissues; S.L. Kloet and E.de Meijer (Leiden Genome Technology Center) for help with 10X Genomics experiments (cell encapsulation, library preparation, single-cell sequencing, pri-mary data mapping, and quality control); the LUMC Flow Cytometry Core Facility, and the LUMC Light and Electron Microscopy Facility. Illustrations were created with BioRender.

## Author contributions

X.C., M.M., S.S., V.V.O. designed the research, analyzed and interpreted results, wrote the manuscript; X.C., F.E.vdH, V.V.O. performed experiments; M.M., S.S. performed scRNA-Seq analysis; K.N. generated MSGN1^mCherry^ hiPSC reporter line; K.A. generated gene targeting constructs for reporter hiPSC lines; X.C., H.M. performed bulk RNA-seq analysis; C.L.M. designed the research and edited the manuscript.

---

**Figure S1.**
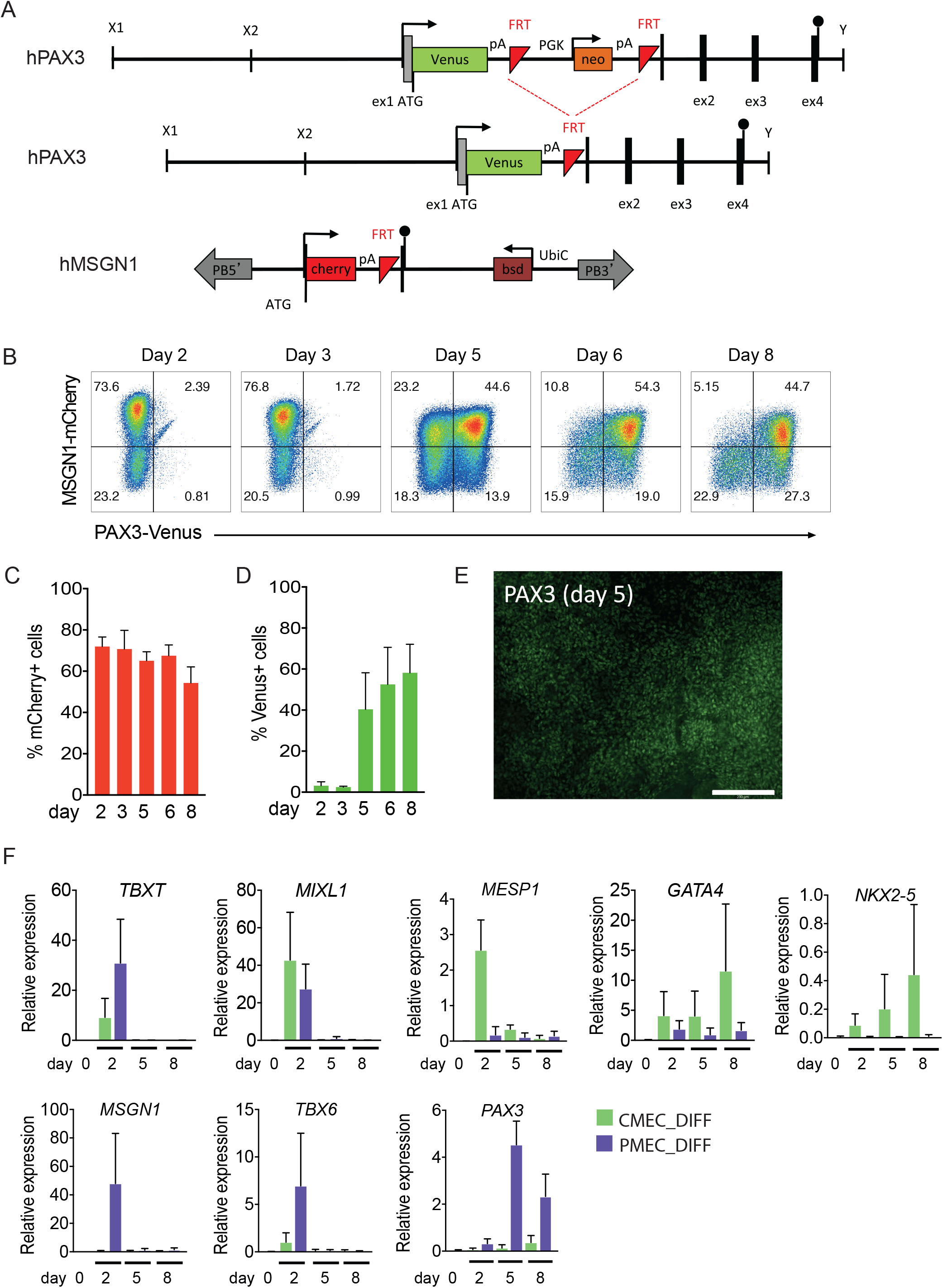
Characterization of PMEC differentiation using MSGN1^mCherry^PAX3^Venus^ dual reporter line. (**A**) Targeting constructs used to generate PAX3-Venus and MSGN1-mCherry hiPSC reporter line. (**B**) Flow cytometry analysis of PAX3^Venus^ and MSGN1^mCherry^ expression on day 2, 3, 5, 6 and 8 of PMEC differentiation. (**C-D**) Quantification of mCherry+ (**C**) and Venus+ (**D**) cells in the total population by flow cytometry on day 2, 3, 5, 6 and 8. (**E**) Representative fluorescence image of PAX3^Venus^ expression on day 5 of PMEC differentiation. Scale bar: 200 μm. (**F**) Quantification of *TBXT, MIXL1, MESP1, GATA4, NKX2-5, MSGN1, TBX6* and *PAX3* expression by qPCR on day 0, 2, 5 and 8 of CMEC (green) and PMEC (purple) differentiation.

**Figure S2.**
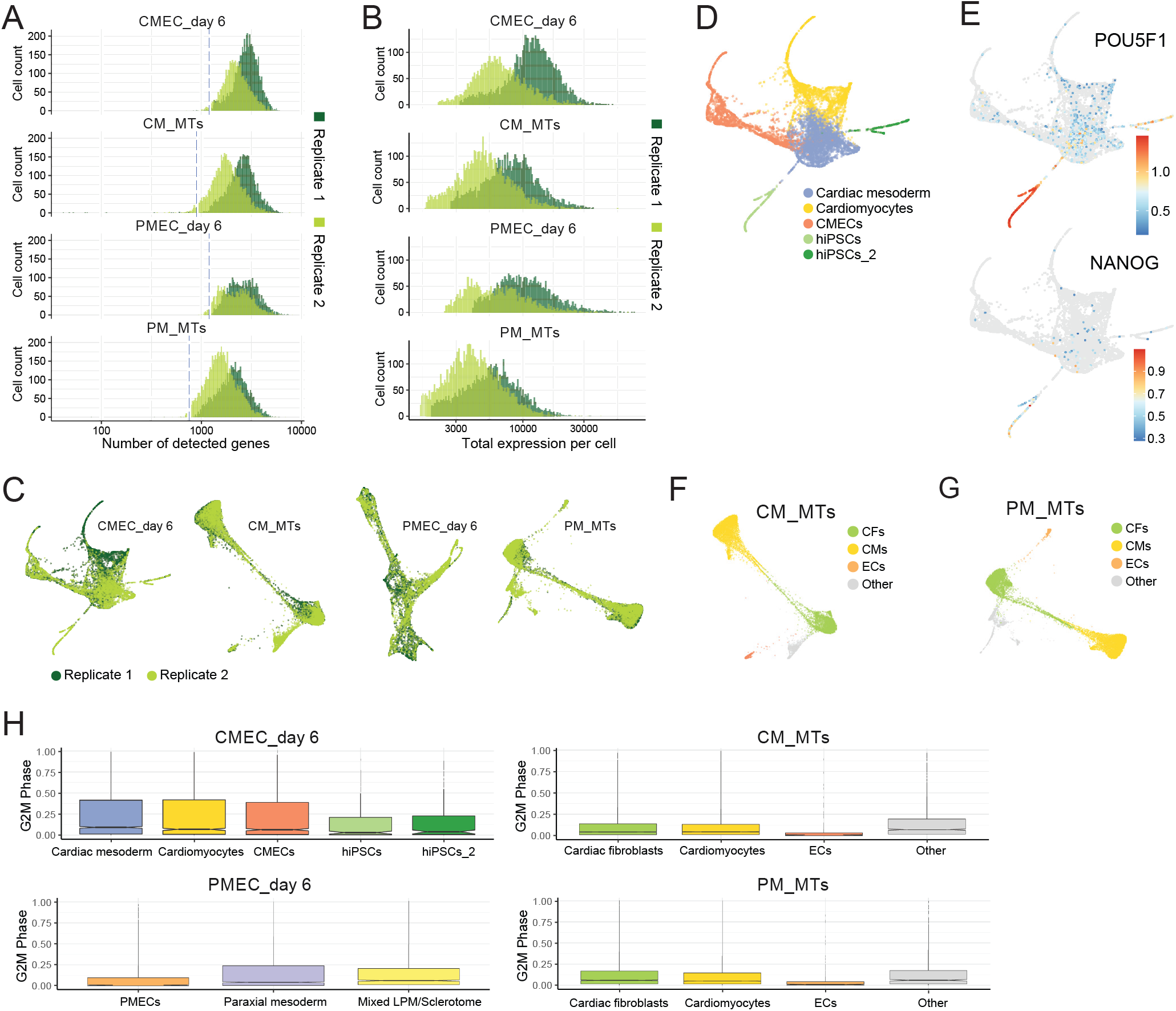
Quality control of scRNAseq datasets. (**A-B**) Distribution of the number of detected genes (**A**) and total expression (**B**) in each cell of the scRNAseq datasets. The dotted blue lines indicate quality control thresholds. Two different batches are labelled with different colors. (**C**) Two different batches of cells collected for each scRNAseq dataset were visualized with PAGA. (**D**) scRNAseq data of CMECs on day 6 is visualized using PAGA. Five cell clusters were identified and labelled with different colors. (**E**) Expression of pluripotency genes *POU5F1* and *NANOG* in the CMECa dataset on day 6 is shown in PAGA plot. Color represents log transformed expression. (**F-G**) scRNAseq data of CM_MTs (**F**) and PM_MTs (**G**) were visualized using PAGA. Four cell clusters were identified. Clusters labelled with “Other” were excluded from downstream analysis. (**H**) Boxplot of G2M phase-score in individual clusters of each dataset.

**Figure S3.**
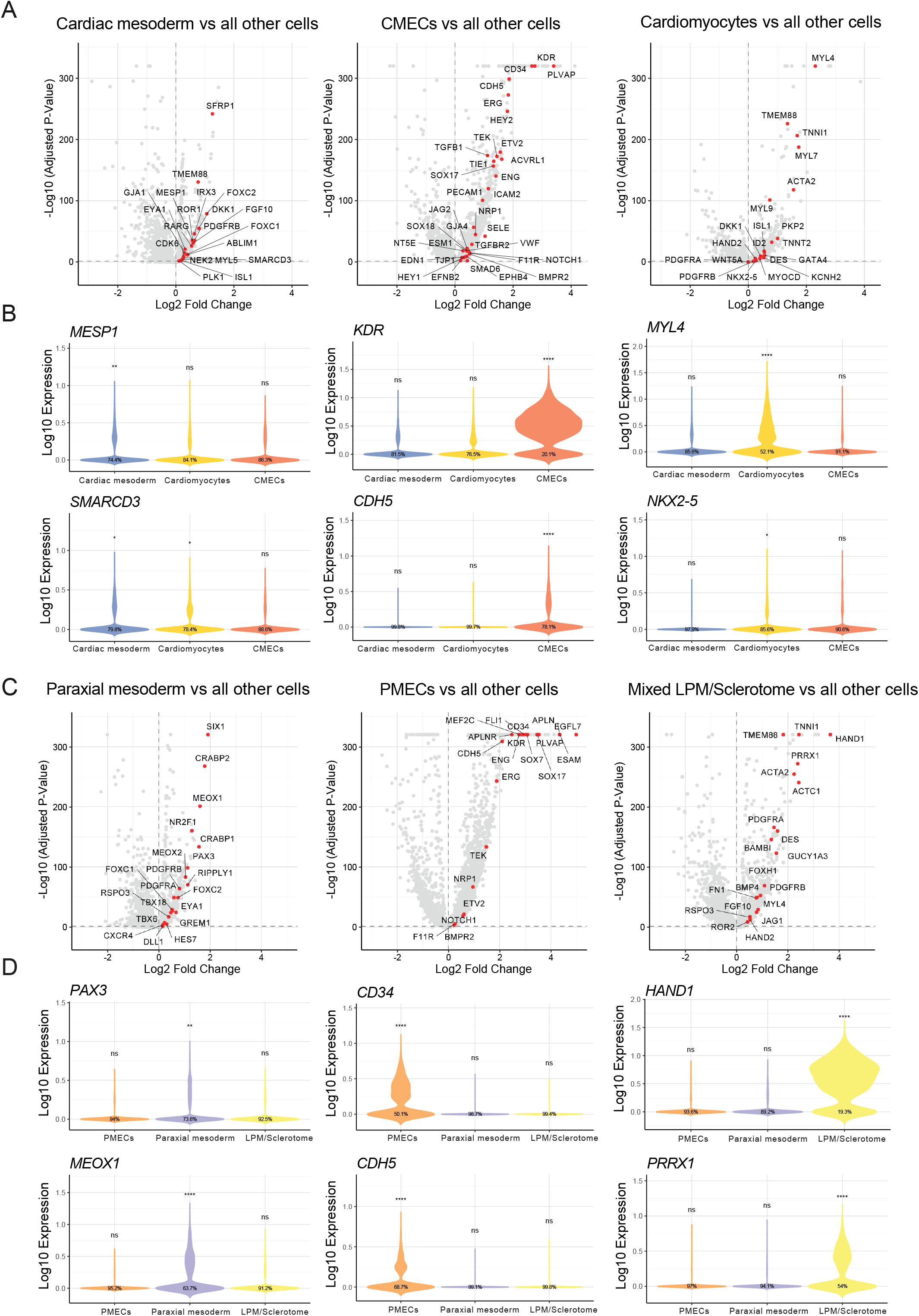
scRNAseq analysis of CMEC and PMEC datasets on day 6. (**A**) Volcano plots showing fold changes and p-values of differential expression tests between individual CMEC clusters and all other cells. Representative significantly up-regulated genes (p_adjusted_ < 0.05 & fold-change > 1.2) are labelled in red. (**B**) *MESP1, SMARCD3, KDR, CDH5, MYL4* and *NKX2-5* expression (log transformed) in three clusters of the CMEC dataset on day 6. (**C**) Volcano plots showing fold-changes and p-values of differential expression tests between individual PMEC clusters and all other cells. Representative significantly up-regulated genes (p_adjusted_<0.05 & fold change > 1.2) are labelled in red. (**D**) *PAX3, MEOX1, CD34, CDH5, HAND1* and *PRRX1* expression (log transformed) in three clusters of PMEC dataset on day 6.

**Figure S4.**
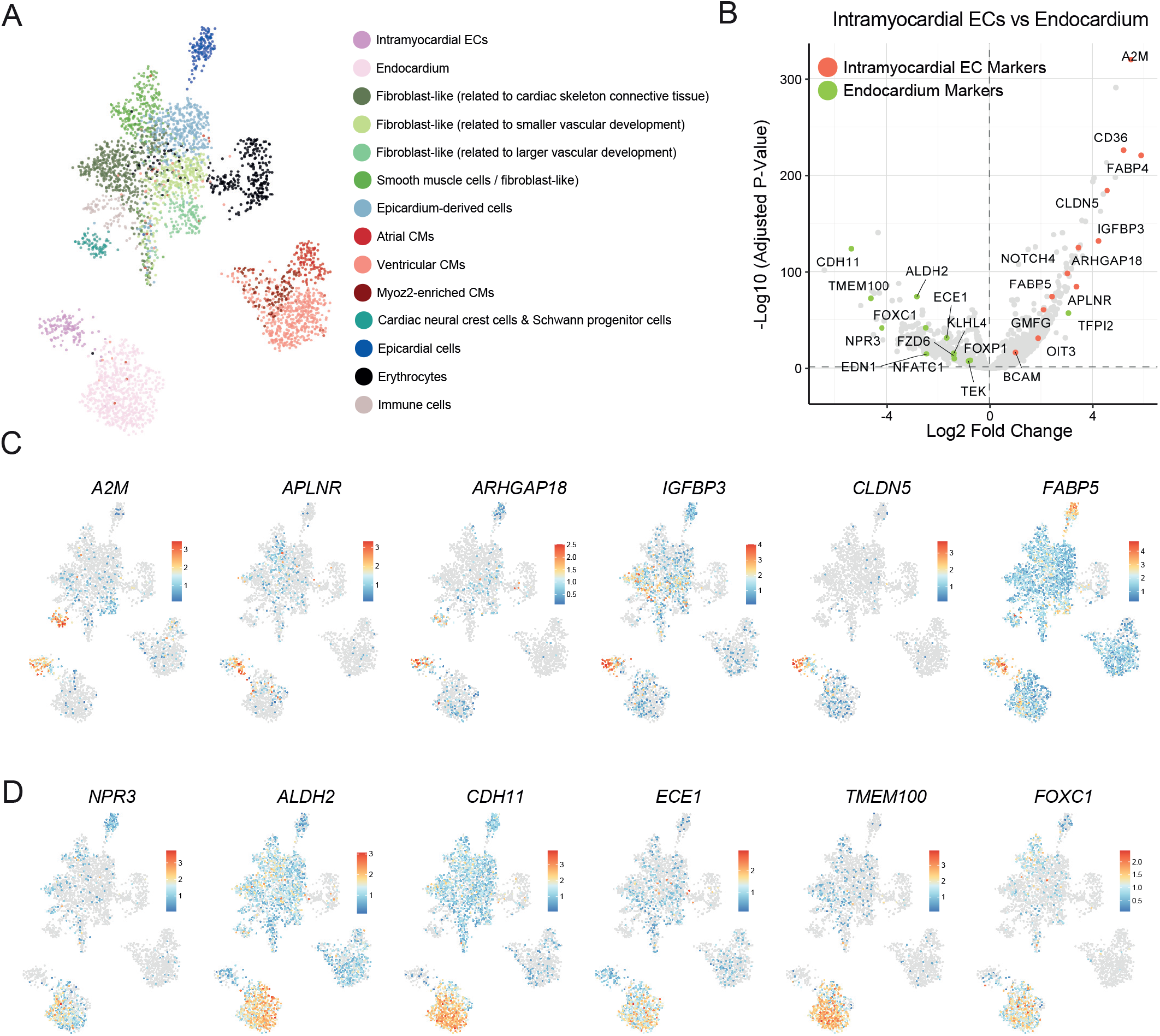
Re-analysis of a published scRNAseq dataset to identify organ specific signatures of human fetal heart ECs. (**A**) Low-dimensional representation (UMAP) of scRNAseq of the human fetal heart (Asp et al., 2019). 14 cell clusters were identified and named based on the original publication, except for two EC clusters: intramyocardial ECs and endocardium. (**B**) Volcano plot showing fold changes and p-values for differential expression tests between intra-myocardial ECs and endocardium in the data set shown in (**A**). Representative differentially expressed genes (p_adjusted_<0.05) that are known as intramyocardial and endocardial markers are labelled in red and green respectively. (**C-D**) Low-dimensional representation (UMAP) of scRNAseq of the human fetal heart (Asp et al., 2019). Log-transformed expression of representative intramyocardial EC markers (**C**) and endocardium markers (**D**) is indicated by color.

**Figure S5.**
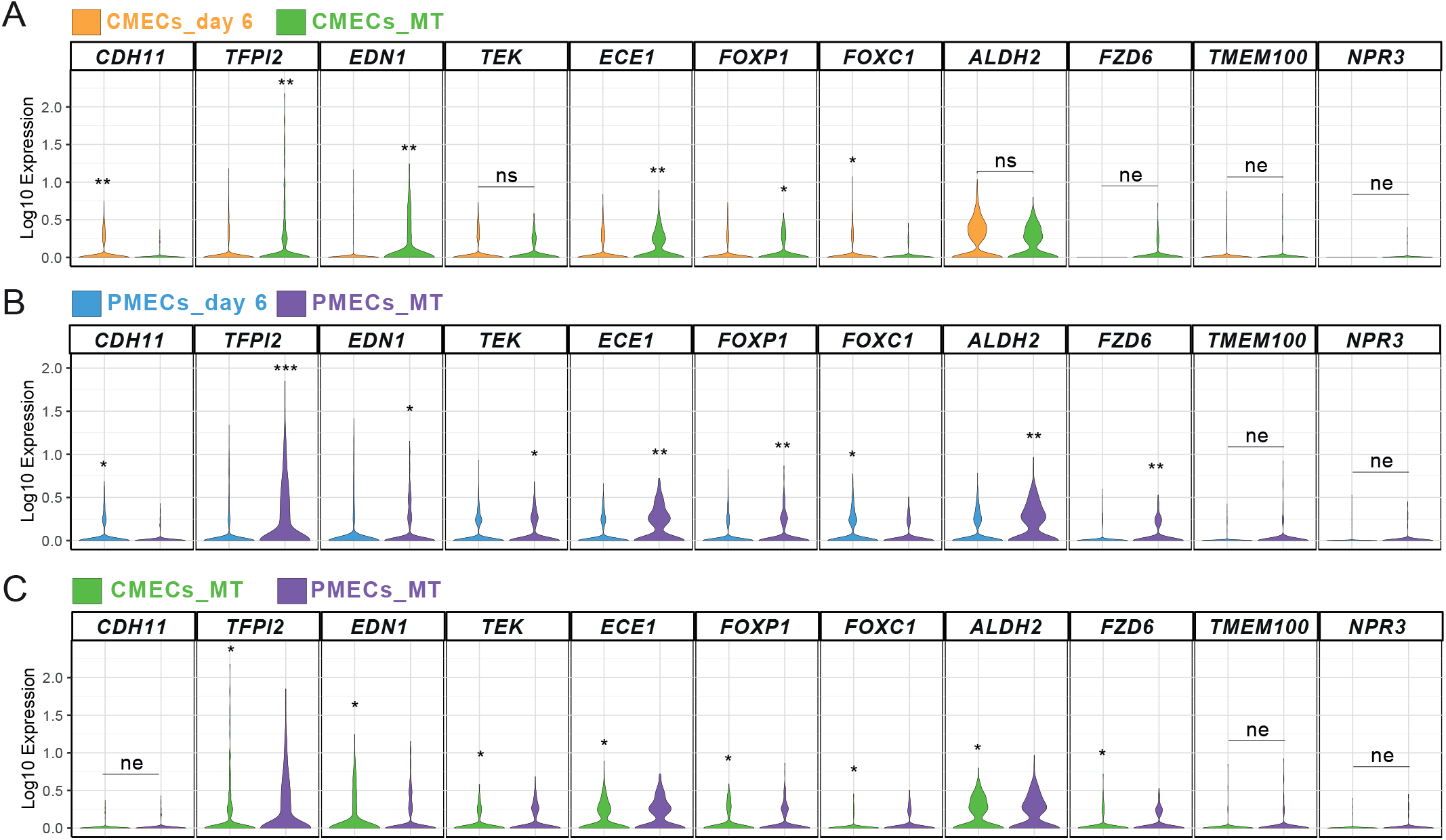
Comparison of organ-specific signatures of hiPSC-ECs on day 6 with ECs in MTs. (**A-C**) Differential expression test between clusters CMECs_day 6 and CMECs_MT (**A**), PMECs_ day 6 and PMECs_MT (**B**), CMECs_MT and PMECs_MT (**C**) for representative endocardial EC markers. ns: p>0.05; * p <=0.05; ** p<= 1e-10; *** p<= 1e-100; **** p<= 1e-200. Clusters with higher expression value were indicated with stars. ne: not expressed (0 counts) in >85% of cells in both groups.

